# Rapid neutralizing assay for circulating H5N1 influenza virus in dairy cows

**DOI:** 10.1101/2024.07.30.605731

**Authors:** Kei Miyakawa, Makoto Ota, Kaori Sano, Fumitaka Momose, Takashi Okura, Noriko Kishida, Tomoko Arita, Yasushi Suzuki, Masayuki Shirakura, Hideki Asanuma, Shinji Watanabe, Akihide Ryo, Hideki Hasegawa

**Author notes:** **Correspondence author:** Kei Miyakawa, Ph.D.

## Abstract

A rapid and safe neutralization assay is required for emerging highly pathogenic avian influenza viruses, including the H5N1 subtype, which was recently found in cows. Herein, we report a novel neutralization assay using HiBiT-tagged virus-like particles (hiVLPs). Our hiVLP-based neutralization test demonstrated a higher quantitative value and shorter assay time than conventional methods. We used this assay to evaluate whether the neutralizing antibodies induced by the candidate vaccine virus (NIID-002) were cross-reactive with cow-derived H5N1. Our results suggest that the circulating H5N1 virus in cows shares antigenic characteristics with NIID-002, providing significant implications for the development and preparation of vaccines.

## Main text

Highly pathogenic avian influenza (HPAI) viruses pose a significant threat to poultry and human health^1,2^. To mitigate these threats, various countries have stockpiled vaccines to prepare for potential pandemics^3,4^. In March 2024, an HPAI (subtype H5N1, clade 2.3.4.4b) was detected in dairy cows in the United States^5^, and spillover to humans was reported^6,7^. Although the World Health Organization (WHO) has designated two candidate vaccine viruses (CVVs) for this clade, i.e., NIID-002 and IDCDC-RG78A^8^, it is critical to rapidly determine the antigenic similarity between the CVVs and HPAI viruses circulating in dairy cows.

The virus neutralization test (VNT) is the gold standard for evaluating vaccine-induced neutralizing antibodies (nAbs). However, this test is time-consuming and requires biosafety level 3 (BSL3) facilities to handle authentic HPAI viruses. To address these limitations, pseudovirus-based VNT (PVNT) has been developed^9,10^; however, a pseudovirus that mimics cow-derived H5N1 has not been previously reported.

We have previously developed a PVNT for SARS-CoV-2 using HiBiT-tagged virus-like particles (hiVLPs)^11^, and therefore sought to apply this technology to influenza viruses. These hiVLPs are composed of the HIV-1 GagPol protein, which forms self-assembling, non-replicating, non-pathogenic entities similar in size and conformation to infectious virions (**Figure 1A**). The HiBiT tag on the viral particles complements the LgBiT protein expressed in target cells, reconstituting a functional luciferase enzyme upon successful viral entry (**Figure 1B**). The resulting luminescence signal is directly proportional to the number of viral particles entering the cells. These characteristics allow the use of BSL2 facilities and rapid quantification of entry into cells expressing the LgBiT protein.

**Figure 1.**
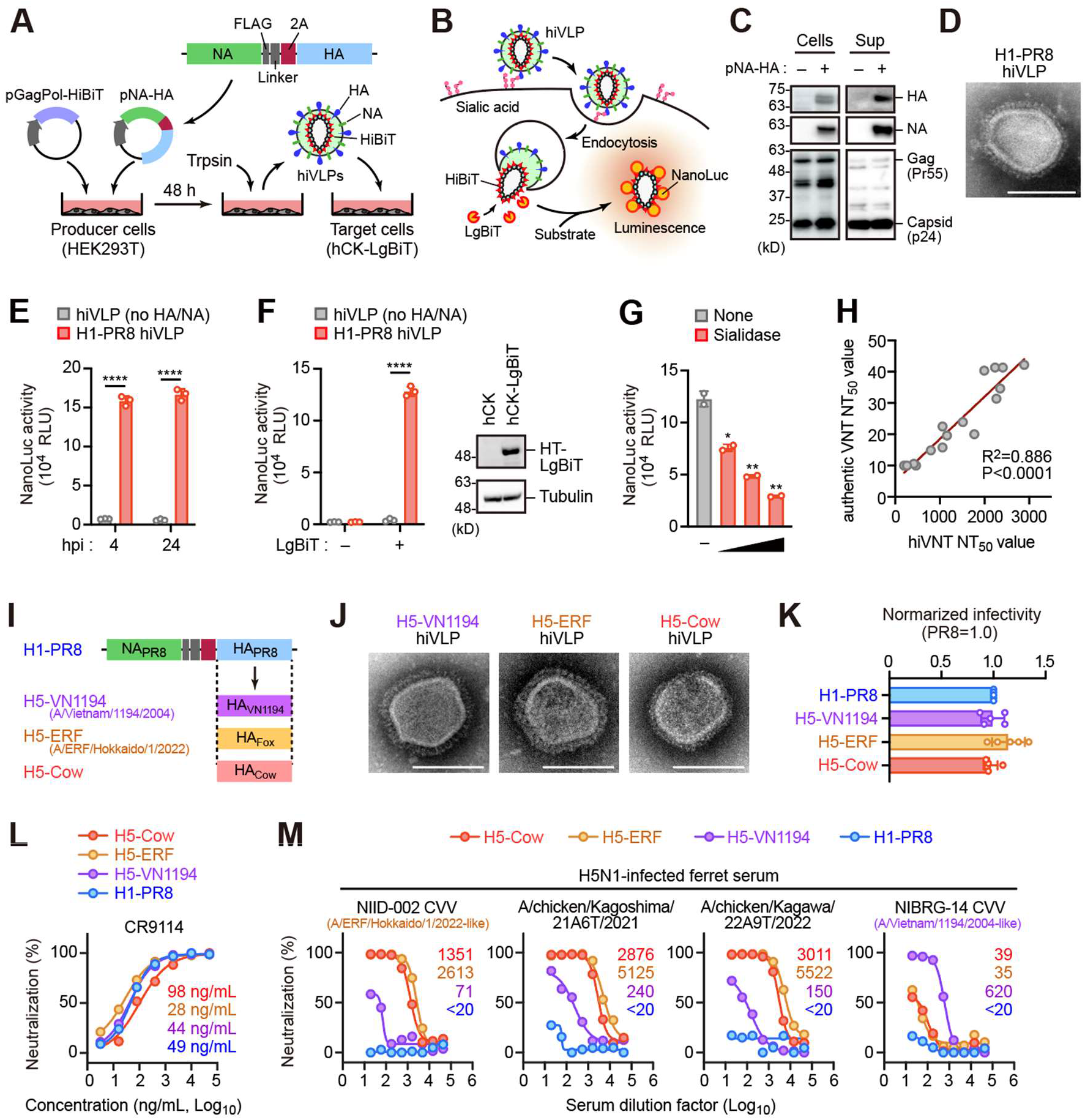
Development of rapid neutralizing test for influenza virus. **(A, B)** Generation of HiBiT-tagged virus-like particles (hiVLPs) mimicking influenza viruses. hiVLPs are produced by co-expressing HIV-1 GagPol genes fused with HiBiT and influenza virus HA and NA glycoprotein genes (pNA-HA) in HEK293T cells (A). The virions emit light upon entry into LgBiT-expressing target hCK cells (B). **(C)** Western blotting of virion-producing cells and their culture supernatants (Sup), probed with anti-HA, NA, and HIV-1 Gag antibodies. **(D)** Electron microscopy image of the H1-PR8 hiVLP. Scale bar, 100 µm. **(E)** Luminescence of hCK-LgBiT cells after 4 and 24 h of indicated hiVLP infection. **(F)** Luminescence in hiVLP-infected hCK cells in the presence and absence of LgBiT. Expression of LgBiT was confirmed through Western blotting. **(G)** hCK-LgBiT cells pre-treated with sialidase (50, 500, and 5000 U/mL) were infected with hiVLP. Cell luminescence was measured 4 h after infection. **(H)** Correlation of 50% neutralizing titer (NT_50_) calculated by conventional virus neutralization test (VNT) and hiVLP-based VNT. **(I)** Construction of pNA-HA plasmid series encoding HA sequence from H5-VN1194 (A/Vietnam/1194/2004), H5-ERF (A/Ezo red fox/Hokkaido/1/2022), and H5-Cow (A/dairy cow/Texas/24-008749-001/2024). Note that NA was equivalent to H1-PR8. **(J)** Electron microscopy image of the hiVLPs shown. Scale bar, 100 µm. **(K)** Normalized infectivity of indicated hiVLPs. Luminescence of hiVLP-infected hCK-LgBiT cells was normalized by particle volume (HiBiT activity). **(L)** Neutralization curve and 50% inhibition concentration (IC_50_) values of CR9114 against indicated hiVLPs. **(M)** Neutralization curves and NT_50_ values for each ferret serum using the indicated hiVLPs.

To generate hiVLPs bearing surface glycoproteins of the influenza virus, HEK293 cells were co-transfected with vectors encoding hemagglutinin (HA), neuraminidase (NA), and HiBiT-fused GagPol (**Figure 1A**). First, we used the HA and NA glycoproteins from the well-studied H1N1 strain (A/Puerto Rico/8/1934). Our experiments revealed that the 2A sequence-linked co-expression of HA and NA resulted in efficient and functional incorporation of both proteins into virions (**Figure 1A**). Subsequently, hiVLPs (designated as H1-PR8) were harvested from HEK293T producer cells, and their expression in cells and virions was confirmed through western blotting (**Figure 1C**) and electron microscopy (**Figure 1D, Supplementary Figure 1A**). Upon the addition of virions to LgBiT-expressing hCK cells^12^, cell luminescence was observed within 4 h and remained stable for up to 24 h (**Figure 1E**). This luminescence was not observed in hCK cells lacking LgBiT expression or in sialidase-treated cells (**Figure 1F, G**). Furthermore, treatment with a clathrin-mediated endocytosis inhibitor (PitStop 2) attenuated the luminescence without observable cytotoxicity (**Supplementary Figure 1B**). These results indicate that H1-PR8 virions can enter cells through mechanisms analogous to those of authentic influenza viruses.

To evaluate the efficacy of hiVLPs as surrogates for authentic viruses in neutralization assays, we measured the 50% neutralization titer (NT_50_) in randomly selected human serum samples using both authentic viruses and hiVLPs. The results revealed a significant correlation between the two methods (**Figure 1H**). Notably, the NT_50_ values calculated with authentic viruses ranged from 10 to 50, whereas those calculated with hiVLPs expanded to a range of 10 to 3000, suggesting that our hiVLP-based PVNT provides a more quantitative assessment of neutralizing activity than conventional VNT.

Subsequently, we adapted the hiVLP-based PVNT to the HPAI virus by substituting the HA proteins in the H1-PR8 hiVLPs with those derived from dairy cows (H5-Cow) and H5N1 CVVs [A/Vietnam/1194/2004 (H5-VN1194) for clade 1 and A/Ezo Red Fox/Hokkaido/1/2022 (H5-ERF) for clade 2.3.4.4b] (**Figure 1I**). We confirmed the formation of virions carrying H5 HA through electron microscopy (**Figure 1J**), and these particles exhibited efficient infectivity (**Figure 1K**). The nAb CR9114, which shows broadly cross-reactivity to influenza viruses^13^, significantly inhibited the cell entry of these virions (**Figure 1L**). Notably, ferret sera immunized with the CVV NIID-002 (clade 2.3.4.4b, A/Ezo Red Fox/Hokkaido/1/2022-like) significantly inhibited infection of H5-Cow and H5-ERF virions (**Figure 1L**). Additionally, two ferret sera against H5N1 clade 2.3.4.4b recently isolated in Japan (A/chicken/Kagoshima/21A6T/2021 and A/chicken/Kagawa/22A9T/2022), also showed good protection against H5-Cow and H5-ERF virions (**Figure 1M**). Conversely, antisera against the CVV NIBRG-14 (clade 1, A/Vietnam/1194/2004-like) exhibited minimal or no protection against the H5-Cow and H5-ERF virions (**Figure 1M**). These results suggest that currently circulating H5N1 virus in dairy cows shares antigenic characteristics with NIID-002 and recently isolated H5N1 clade 2.3.4.4b viruses.

This study had several limitations. First, the H5N1 hiVLPs reported in this study were generated by replacing only HA with the H5 subtype. Second, the M2 glycoprotein was not incorporated into the virions. Therefore, our system is unsuitable for the detection of nAbs targeting NA or M2. We are currently improving hiVLPs to detect nAbs against these antigens. Nonetheless, given that most broadly nAbs bind to the highly conserved stem region of HA among influenza virus subtypes^14,15^, hiVLP-based PVNT represents a potential alternative to conventional VNT. This novel approach has significant implications for the development and preparation of vaccines.

## Acknowledgements

The authors thank Kenji Yoshihara for providing technical assistance. The authors also thank Dr. Othmar Engelhardt (Medicines and Healthcare products Regulatory Agency), Dr. Yoshihiro Sakoda (Hokkaido University), and Dr. Yuko Uchida (National Agriculture and Food Research Organization) for providing H5N1 viruses, Dr. Yoshihiro Kawaoka (University of Wisconsin) for providing hCK cells, and Dr. Yuko Morikawa (Kitasato University) for providing the reagents.

