## Supporting information for "Rapid neutralizing assay for circulating H5N1 influenza virus in dairy cows"

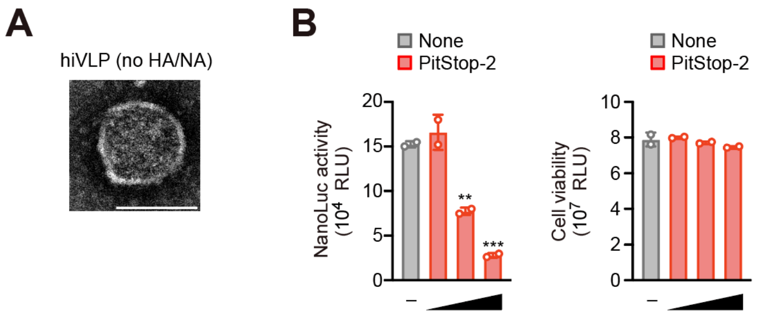


**Supplementary Figure 1. Characteristics of the hiVLPs.**

**(A)** Electron microscopy image of the hiVLP not expressing HA and NA. Scale bar, 100 µm.

**(B)** hCK-LgBiT cells pre-treated with PitStop 2 (20, 40, and 80 µM) were infected with hiVLP. Cell luminescence was measured 4 h after infection (left). Cell viability was also shown (right).

### Supplementary Materials and Methods

#### Cells

HEK293T (ATCC, #CRL-3216) and hCK [1] (kindly provided by Prof. Yoshihiro Kawaoka) cells were maintained in Dulbecco's Modified Eagle Medium (DMEM) supplemented with 10% fetal bovine serum (FBS). hCK-LgBiT cells were engineered by lentiviral transduction with a vector encoding HaloTag-LgBiT and subsequently selected using blasticidin S (10 μg/mL).

#### Plasmids

The pGagPol-HiBiT plasmid has been previously described [2]. We constructed NA-HA bicistronic plasmids (pNA-HA series) based on pCAHyg-NA2aHA-4d vector. The open reading frame of pCAHyg-NA2aHA-4d comprises the full-length NA sequence (A/PR8), a FLAG epitope, a linker sequence (GSGSG), the 2A-like self-processing sequence of *Thosea asigna* virus, and the full-length HA sequence (A/PR8). Based on the pCAHyg-NA2aHA-4d vector, derivatives were generated by replacing the human codon-optimized HA sequence with that derived from:

A/dairy cow/Texas/24-008749-001/2024 (GISAID #EPI_ISL_19014384)
A/Ezo red fox/Hokkaido/1/2022 (GISAID #EPI_ISL_12174842)
A/Vietnam/1194/2004 (NCBI GenBank #AAT73273.1)

#### Viruses

A/chicken/Kagoshima/21A6T/2021 (H5N1; GISAID #EPI_ISL_6829533) and A/chicken/Kagawa/22A9T/2022 (H5N1; GISAID #EPI_ISL_18286296) were provided by the National Agriculture and Food Research Organization, Japan. NIBRG-14 (A/Vietnam/1194/2004; GISAID #EPI_ISL_314984) was provided by the Medicines and Healthcare products Regulatory Agency, UK. NIID-002 (GenBank #LC831696, LC831697) possessing the HA and NA genes from A/Ezo red fox/Hokkaido/1/2022 (H5N1; GISAID #EPI_ISL_12174842), and six internal genes from A/Puerto Rico/8/34 was generated by reverse genetics [3]. These viruses were propagated in the allantoic cavities of 10-day-old embryonated chicken eggs and virus titers were determined by the 50% egg infectious dose ($EID_{50}$).

#### Western blotting

Protein samples were prepared in sodium dodecyl sulfate (SDS) loading buffer and separated on 10-20% gradient polyacrylamide gels (Wako). Electrophoresis was performed under standard conditions, followed by electroblotting onto polyvinylidene difluoride (PVDF) membranes (Merck). The membranes were probed with anti-H1N1 HA (Genetex, #GTX127357), anti-H1N1 NA (Genetex, #GTX629696), or anti-HIV p24 (clone 183-H12-5C, NIH AIDS Reagent Program #3537) antibodies and horseradish peroxidase-conjugated secondary antibodies (Cytiva). Protein bands were visualized using the ImageQuant 800 (Cytiva), according to the manufacturer's instructions.

**Production of hiVLPs**

HEK293T cells were co-transfected with pHIV-GagPol-HiBiT and pNA-HA at a ratio of 1:1 using the Lipofectamine 3000 transfection reagent (Thermo Fisher Scientific). Approximately 4 h post-transfection, the culture medium was replaced with Opti-MEM reduced-serum medium to optimize hiVLP formation. At 48 h post-transfection, acetylated trypsin (Sigma) was added to the culture supernatant at a final concentration of 3 µg/mL and incubated for 30 min at 37 °C. This step ensures proper cleavage and activation of the hemagglutinin protein. Subsequently, the supernatant containing hiVLPs was harvested and filtered through a 0.45-µm Millex-HV filter (Merck) to remove cellular debris.

**Neutralizing assay using hiVLPs**

Prior to infection, 50 µL of DMEM containing hiVLPs (approximately $10^7$ relative light units (RLU) of HiBiT activity) and serially diluted serum (1:20–1:43,740 dilution) or neutralizing antibody (3.2–50,000 ng/mL) were incubated for 30 min at 37 ºC. hCK-LgBiT cells seeded in 96-well plates were inoculated with the pre-incubated hiVLP solution for 4 or 24 h at 37 °C in a 5% $CO_2$ humidified incubator. For several experiments, hCK-LgBiT cells were pre-treated with α2-3,6,8 neuraminidase (NEB; 50, 500, and 5,000 U/mL) or Pitstop 2 (MedChemExpress; 20, 40, and 80 µM) for 30 min prior to infection. Following the infection period, cells were then washed with phosphate-buffered saline (PBS) and treated with 100 µL of PBS containing 1 µM DrkBiT peptide (Promega) for 2 min at room temperature. Subsequently, 25 µL of Nano-Glo Live Cell Substrate (Promega) was added to each well. Luciferase activity was measured using a GloMax Discover System (Promega). The 50% neutralizing titer ($NT_{50}$) was calculated based on the half-maximal inhibitory concentrations determined using the Prism 10.2.2 software (GraphPad).

**Neutralizing assay using authentic viruses**

A 50 µL mixture of DMEM containing influenza virus (A/Puerto Rico/8/1934, 100 $TCID_{50}$) and serially diluted human serum (1:20–1:43,740 dilution) was incubated for 30 minutes at 37°C. hCK cells seeded in 96-well plates were inoculated with the pre-incubated virus solution for 1 h at 37 °C in a 5% $CO_2$ humidified incubator. Following infection, 200 µL of DMEM containing acetylated trypsin (3 µg/mL) was added to each well to facilitate multi-cycle viral replication. At 48 h after infection, cells were washed with PBS, and 40 µL of CellTiter-Glo Substrate (Promega) was added to each well, and cell viability was measured using a GloMax Discover System (Promega). In the absence of neutralizing antibodies, viral infections lead to cell death, which results in decreased luminescence. $NT_{50}$ was calculated based on the half-maximal inhibitory concentrations determined using Prism 10.2.2 software (GraphPad).

**Electron Microscopy**

The hiVLP-containing supernatant was concentrated approximately 80-fold by centrifugation at 10,000 × g for 90 min at 4 °C, and the resulting virion pellet was resuspended in 20 µL of PBS. Approximately 5 µL of concentrated hiVLP solution was applied to carbon-coated copper grids and incubated at room temperature for 1 min. Excess fluid was removed using filter paper, followed by staining with 2% phosphotungstic acid solution for 30 s. The samples were observed using a transmission electron microscope (JEM-1400Plus; JEOL) at an acceleration voltage of 100 kV. The images were captured using a CCD camera (EM-14830RUBY2, JEOL). All of the operations were performed in a clean environment to prevent contamination.

**Ferret immunization and serum collection**

Male ferrets (Mustela putorius furo), 12- to 14-months old and seronegative for the influenza A virus, were obtained from Japan SLC, Inc., Shizuoka, Japan. Ferrets (n=4 per group) were anesthetized intramuscularly with ketamine (25 mg/kg body weight) and xylazine (2 mg/kg body weight) and intranasally inoculated with $10^6$ $EID_{50}$ of NIBRG-14 and NIID-002 and $10^3$ or $10^5$ $EID_{50}$ of Ck/Kagoshima and Ck/Kagawa viruses diluted in 0.5 mL of PBS. Fourteen days post-inoculation, the ferrets received a booster immunization with a mixture of concentrated viruses and adjuvant (Sigma adjuvant system, Sigma). Ferrets inoculated with Ck/Kagoshima and Ck/Kagawa were orally treated for five days post-inoculation with oseltamivir (10 mg/kg of body weight/day) to reduce the symptoms. The ferrets were monitored daily for clinical signs and body weight loss. Any ferret that lost ≥25% of their initial body weight or displayed severe signs of neurological symptoms was humanely euthanized. Seven days after the booster, the ferrets were euthanized by cardiac puncture, and blood samples were collected. Blood was allowed to clot at room temperature for 30 min, and subsequently centrifuged at 2,000 × *g* for 10 min at 4 °C. The serum was diluted with the prepared RDE solution (Denka) at a ratio of 1:3 (v/v). The serum-RDE mixture was incubated at 37 °C for 18 h. Following incubation, the mixture was heat-inactivated at 56 °C for 30 min to denature the RDE. The treated samples were then cooled and diluted with 6× the volume of serum in PBS. Processed sera were aliquoted and stored at -20 °C until further use. All animal procedures were approved by the Institutional Animal Care and Use Committee and were conducted in accordance with the guidelines for animal experiments (approval numbers: 119055, 113135, and 123054).

**Collection of human serum samples**

Serum samples were randomly collected from healthy volunteers between January 2014 and December 2015. All samples were anonymized and obtained from Yokohama City University Biobank. The Serum samples were treated with RDE solution as described above. The study was conducted in accordance with the Declaration of Helsinki and the Ethical Guidelines for Medical Research Involving Human Subjects and was approved by the Yokohama City University Ethics Committee (approval number: B160800009).

**Statistical analysis**

All bar graphs represent mean and standard deviation. The statistical significance of the differences between the two groups was evaluated using a two-tailed unpaired t-test in Prism 10.2.2 software (GraphPad).
